# Inbreeding depression is greater in benign than in stressful environments

**DOI:** 10.64898/2026.07.09.737435

**Authors:** Yuan Fu Chan, Raj Whitlock

**Author notes:** **Competing Interest Statement:** None.

## Abstract

The potential for environmental change to compound the detrimental effects of inbreeding depression in small and isolated populations is a significant concern in conservation biology. Previous evidence syntheses suggested that environmental stress exacerbates inbreeding depression, but were based on limited data. Here, we comprehensively test the relationship between inbreeding depression and environmental stress in natural populations using Bayesian mixed-effects meta-analysis on a large, high-quality data set of 2127 inbreeding depression effect sizes from animals and plants. Our results show that inbreeding depression is significantly higher in benign than in stressful environments. Analyses of both inbreeding depression and stress-induced changes in genetic load supported a unimodal (humped) relationship between the costs of inbreeding and stress intensity, with a peak at intermediate stress. At the highest levels of stress there was, on average, a significantly greater inbreeding load in benign than in stressful environments. We suggest that the lower cost of inbreeding associated with extreme stress results from constraints on the expression of inbreeding depression as fitness and phenotypes decline towards zero. Our findings help to resolve long-standing uncertainty around how inbreeding and environmental change interact, revealing that inbreeding responses vary non-linearly with environmental stress intensity, but showing that stress does not generally amplify inbreeding depression. As such, they will inform both the management of populations of conservation concern and predictions of species’ responses to global environmental change.

## 1 INTRODUCTION

Anthropogenic land and sea use changes have driven widespread habitat fragmentation, isolating and reducing the size of populations of plants and animals (1), and exposing them to inbreeding depression—genetic fitness costs expressed in inbred offspring (2–4). Inbreeding depression is a widespread phenomenon (5) and leads to immediate fitness costs that are sufficiently strong to imperil population viability, requiring the resumption of gene flow to restore individual fitness and demographic stability (6). Small population sizes also lead to the erosion of genetic diversity, limiting evolutionary potential and the opportunity for adaptation to global environmental change in natural populations (7, 8). Inevitably, the demographic changes that potentiate inbreeding depression and the loss of genetic variation will be accompanied by exposure to stress from other global change drivers, including climate change, pollution, and pathogens (3, 9). This begs the questions of whether stress induced by environmental change will compound fitness costs associated with inbreeding depression, raising an immediate challenge to demographic sustainability, and in the longer term, whether accompanying genetic erosion will preclude the capacity for genetic rescue via adaptation. As a result, understanding how environmental changes modify the effects of inbreeding depression is a pressing issue for the management of populations of conservation concern.

The extent and expression of inbreeding depression can be highly dependent on the environmental conditions under which it is being measured, often through specific forms of genotype-by-environment interaction; a phenomenon known as environment-dependent inbreeding depression (10–13). Both increases and decreases in inbreeding depression have been observed with increased environmental stress. In 2010, Fox and Reed reported that inbreeding depression increased under stressful conditions in the seed-feeding beetle *Callosobruchus maculatus*. Similarly, Kristensen *et al*. (15) demonstrated that inbreeding depression in egg-to-adult viability of *Drosophila melanogaster* occurred only under stressful temperature regimes. In contrast, Henry and colleagues (12) found lower inbreeding depression in survival under field compared with lab conditions in the freshwater snail, *Physa acuta* (field conditions were assumed to be more stressful). Likewise, Sandner and Matthies (16) showed that inbreeding depression in *Silene vulgaris* was less severe under nutrient deficiency and drought stress.

Environment-dependent inbreeding depression arises via two key mechanisms. The first involves changes in phenotypic differences between inbred and outbred offspring over environmental gradients, known as environment-dependent phenotypic expression (10). The second comprises environmental changes in selection regimes that alter the costs of inbreeding, referred to as environment-dependent selection (10, 17, 18). Under environment-dependent phenotypic expression, the environment shapes the expression of phenotypes differently for outbred and inbred individuals, under a constant regime of natural selection, either through direct plastic changes in phenotypes or gene expression, or indirectly by modifying stress tolerance or physiology (10, 19–21). To do so, environmental change must alter the phenotypic effects of the detrimental alleles underpinning inbreeding depression. The genetic basis of inbreeding depression (i.e. the number and identity of contributing alleles) is constant. Inbred and outbred progeny thus respond to environmental factors via distinct reaction norms altering fitness differences between them in different environments.

Under the second mechanism, the environment does not modify the expression of phenotypes, but alters selection such that fitness differences between inbred and outbred progeny are environmentally contingent (10). Specifically, this mechanism implies that changing conditions modify the selection coefficients of the alleles underlying inbreeding depression and thereby vary their fitness consequences without altering phenotypes. This could involve either changes in the average negative effects of the detrimental alleles underpinning genetic load (22, 23) or shifts in their identity and number, as previously neutral or beneficial alleles become detrimental (24–26). The two mechanisms that drive environment-dependent inbreeding depression are not mutually exclusive. For example, a thermal shift in inbreeding depression could imply both changes in selection regime on growth rate and phenotypic shifts in development time.

Several evidence syntheses have addressed the relationship between environmental stress and inbreeding depression, seeking a general pattern. A pioneering study conducted by Armbruster and Reed (27), compared genetic load under benign and stressful environments, across 34 studies. They found that inbreeding costs were significantly greater in stressful environments, demonstrating environment-dependent inbreeding depression. Later, Fox and Reed (14) performed a meta-analysis quantifying the change in lethal equivalents between stressful and benign environments over a gradient of environmental stress. This approach permitted an evaluation of stress-induced changes in inbreeding load within populations in relation to the intensity of stress. Their results indicated that the number of lethal equivalents increases strongly and linearly with the intensity of the environmental stress. However, their dataset was small, and contained few data points at high stress levels. Thus, it was not clear whether stress-induced shifts in inbreeding depression should vary linearly with environmental stress intensity.

Individual empirical studies have suggested that the relationship between inbreeding depression and environmental stress is nonlinear. Using populations of *Drosophila melanogaster*, Schou et al., (28) found the rate of increase in inbreeding depression for egg-to-adult viability decreased at higher stress levels. Likewise, Springer et al., (29) reported a lack of inbreeding-stress interaction in *C. maculatus* under severe stress when comparing their study with Fox and Reed’s (14). They suggested that this discrepancy arose from differences in the intensity of stress applied across studies: in their experiment, severe stress substantially reduced overall fitness, thereby limiting the capacity for inbreeding depression to be expressed. This nonlinearity occurs because at extreme stress levels, by definition, fitness in outbred individuals is very low (approaching zero) relative to that in benign environments, constraining the potential for inbreeding depression (29, 30). In addition, there are limits on the extent to which the fitness costs associated with individual genotypes can increase: once a genotype becomes lethal, more extreme detrimental effects are not possible. Together, these constraints imply a unimodal relationship between environmental stress and the intensity of inbreeding depression: beyond a point, further increases in stress will no longer amplify inbreeding depression, and eventually the extent of stress should cause inbreeding depression to be lower than that seen in benign environments.

Using a comprehensive meta-analysis, we evaluate how inbreeding depression in natural populations is modified by environmental stress. First, we determine how inbreeding depression responds, on average, to environmental stress, and how it co-varies with stress intensity. Second, we evaluate the change in inbreeding load between benign and stressful environments, and its relationship with stress intensity. We also investigate variation in inbreeding responses among different types of environmental factors, among taxa and between fitness component and other traits. We predicted that inbreeding depression should increase with environmental stress up to a peak at intermediate stress levels, driven by changes in either phenotypic expression, or the trait–fitness mapping (10, 14). However, beyond this point (i.e. at high stress levels), we predicted that stress-induced shifts in inbreeding depression should decrease, limited by constraints to the expression of inbreeding depression in stressful environments.

## 2 RESULTS

Our meta-analysis captured data from 116 articles, with a total of 2127 effect sizes, describing how inbreeding depression in natural populations changes under environmental stress. This evidence base included animals (39.1% of data) and plants (60.9%) with most studies from Europe, and North America. We detected significant inbreeding depression in both fitness components (survival, reproduction, viability, fitness) and other (non-fitness component) traits in both benign and stressful environments (linear mixed-effects meta-analysis; *P* < 0.001). On average, inbreeding depression was significantly lower in stressful environments than benign environments across populations and taxa (*P* = 0.002; Fig. 1). Inbreeding depression was also substantially and significantly greater (by 19%) in fitness-component traits than non-fitness component traits (*P* = 0.032; Fig. 1). The presence of environmental stress did not modify variation in inbreeding depression among these trait classes (stress × trait class interaction; *P* > 0.05).

**Fig. 1.**
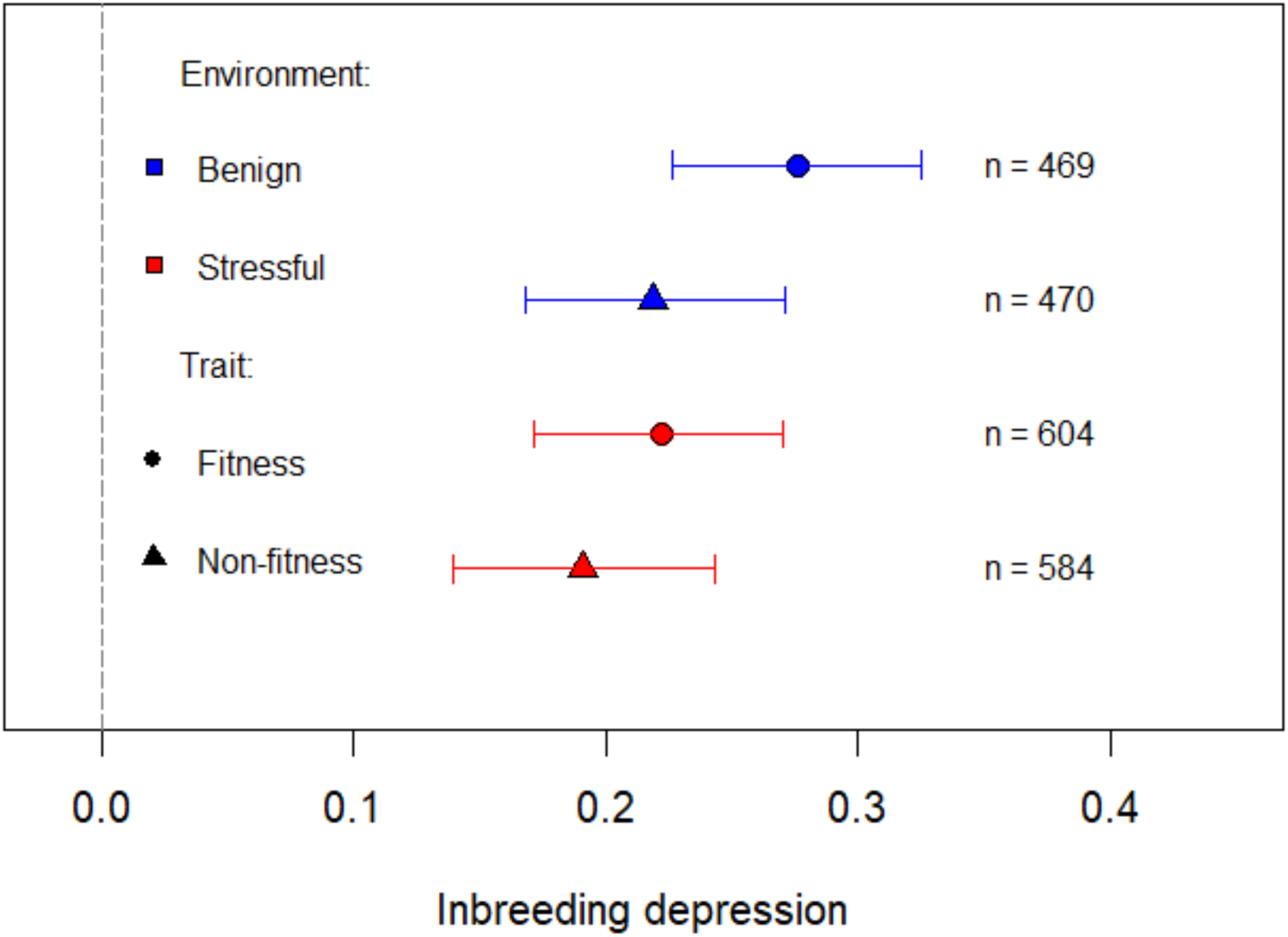
Variation in inbreeding depression between fitness and non-fitness component traits in benign and stressful environments. Positive values indicate inbreeding depression (a cost to fitness or phenotype). Points and error bars (95% credible intervals) show model predictions based on a model that incorporated a trait class by environment interaction, at an inbreeding coefficient of 0.5; n indicates the number of effect sizes.

### 2.1 Inbreeding depression and environmental stress intensity

Stress intensity was quantified from phenotypic differences among environments in outbred individuals (14). We detected a significant second-order (humped) polynomial relationship between stress intensity and inbreeding depression in fitness components (*P* < 0.001; Fig. 2a; *SI Appendix*; Table S4). On average, inbreeding depression increased with environmental stress, reaching a peak at intermediate stress levels (stress = 0.46). Beyond this point, the magnitude of inbreeding depression decreased (Fig. 2a). In stressful environments, inbreeding depression was therefore most acute at intermediate stress levels. For non-fitness component traits, there was no consistent relationship between stress intensity and inbreeding depression (Slope = 0.059, *P* = 0.41; Fig. 2b). Neither was there a significant relationship between the inbreeding coefficient and inbreeding effect sizes for these traits (*P* = 0.256).

**Fig. 2.**
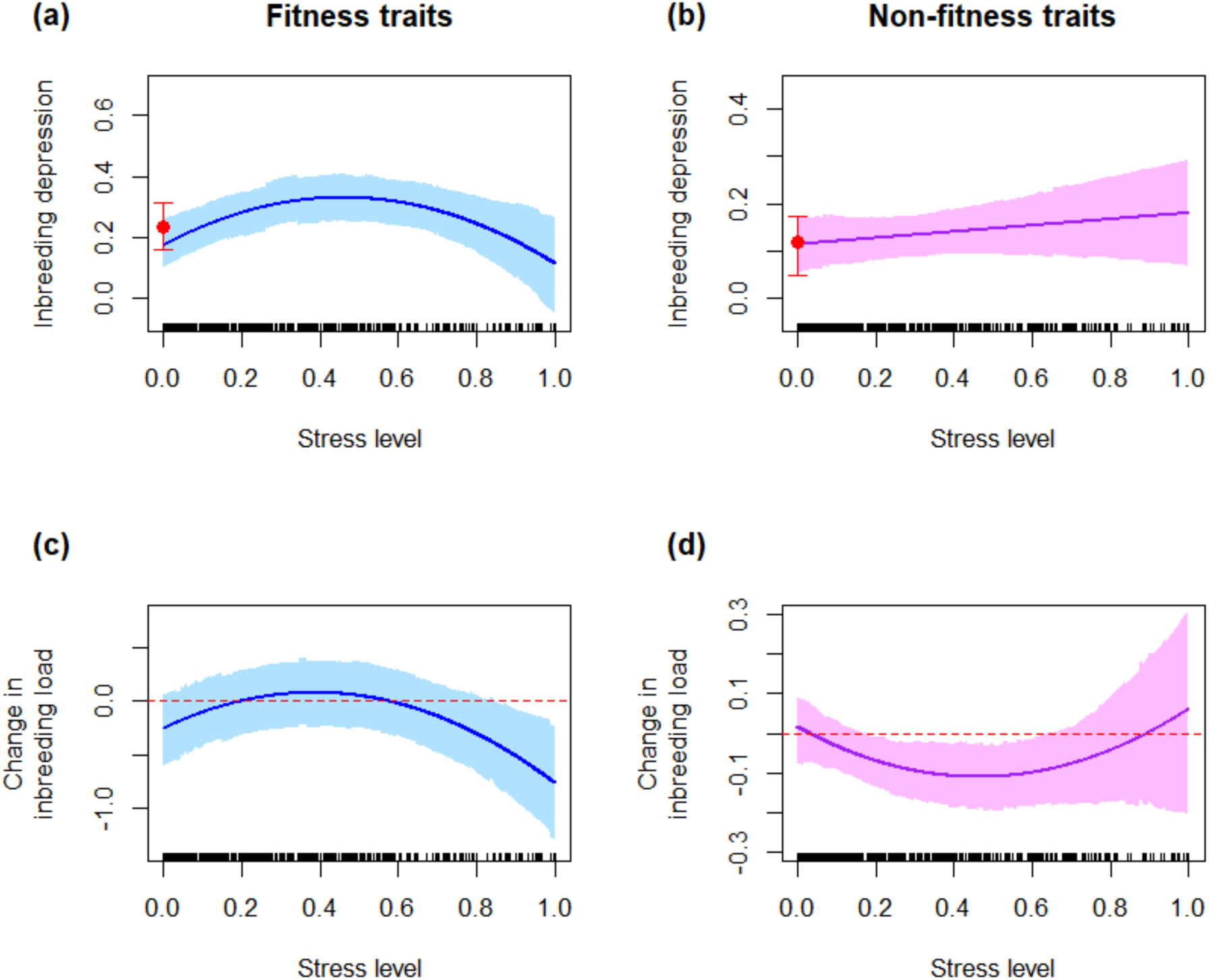
Relationships between environmental stress levels and the expression of inbreeding depression. Each plot shows model predictions for variation of inbreeding depression at an inbreeding coefficient of 0.5 (a) and (b) or the change in inbreeding load between benign and stressful environments (c), (d) with environmental stress level. Stress level data are shown as an x-axis rug. The red points and whiskers show the mean inbreeding effect sizes in benign environments and their 95% credible intervals (undefined for (c) and (d)). (a) the relationship between inbreeding effect sizes and stress level in stressful environments in fitness component traits (second-order polynomial effect shown; *n* = 604 effect sizes). (b) the relationship between inbreeding effect sizes and stress level in stressful environments in non-fitness component traits (linear effect shown; *n* = 584). (c) relationship between stress-induced changes in inbreeding load and stress levels for fitness component traits (second-order polynomial effect shown; *n* = 602). (d) relationship between stress-induced changes in inbreeding load and stress levels for non-fitness traits (linear effect shown; *n* = 584). The shaded areas represent 95% credible zones for the regression line. The red dashed line is y = 0 in (c) and (d).

We calculated the difference in inbreeding load between stressful and benign environments to examine how population-level stress-linked shifts in inbreeding depression varied with stress intensity. On average, stressful environments had a lower inbreeding load than benign environments. This difference was not significantly different from zero for fitness component traits (*P* = 0.536) but had weak statistical support for non-fitness component traits (*P* = 0.082). Mirroring our analysis of the absolute magnitude of inbreeding depression, we detected a significant second-order polynomial relationship between these stress-induced shifts in inbreeding depression and stress intensity for fitness component traits (*P* < 0.001; Fig. 2c; *SI Appendix*; Table S5). At an intermediate stress level (Stress = 0.5), the posterior probability that inbreeding load was stronger in stressful than benign environments was 0.607. This model prediction was consistent with the empirical distribution of observed effect sizes in the stress level range 0.35–0.65, where the proportion of significantly negative changes in inbreeding load in response to stress (benign environments with greater inbreeding load) was comparable with that for significantly positive cases (*SI Appendix*; Table S6). Stress-induced shifts in inbreeding load did not differ from zero for stress levels below 0.852 (Fig. 2c). At higher stress levels than this, inbreeding load in benign environments was on average, significantly stronger than in stressful environments. For non-fitness component traits, we found a weak positive polynomial relationship between stress-linked changes in inbreeding load and the level of environmental stress (Fig. 2d; *P* = 0.078). At intermediate stress, load was stronger in benign environments with a posterior probability of 0.998, but at extreme stress levels, the shift in inbreeding load was not significantly different from zero (*P* = 0.578).

### 2.2 Inbreeding depression under different environmental stress types

We detected significant inbreeding depression under all types of environmental stress except salinity (Fig. 3a). However, there was no significant variation in inbreeding depression between different environments (i.e. the environment predictor did not improve model fit; *SI Appendix*: Table S4). Neither was there significant variation in inbreeding responses between high-level categories of environmental stress (abiotic, biotic or “multiple factors”; Fig 3c). Variation in the change in inbreeding load induced by stress could be explained by neither different types of environments (*SI Appendix*; Table S5; Fig. 3b) nor by high-level environmental categories (Fig. 3d).

**Fig. 3.**
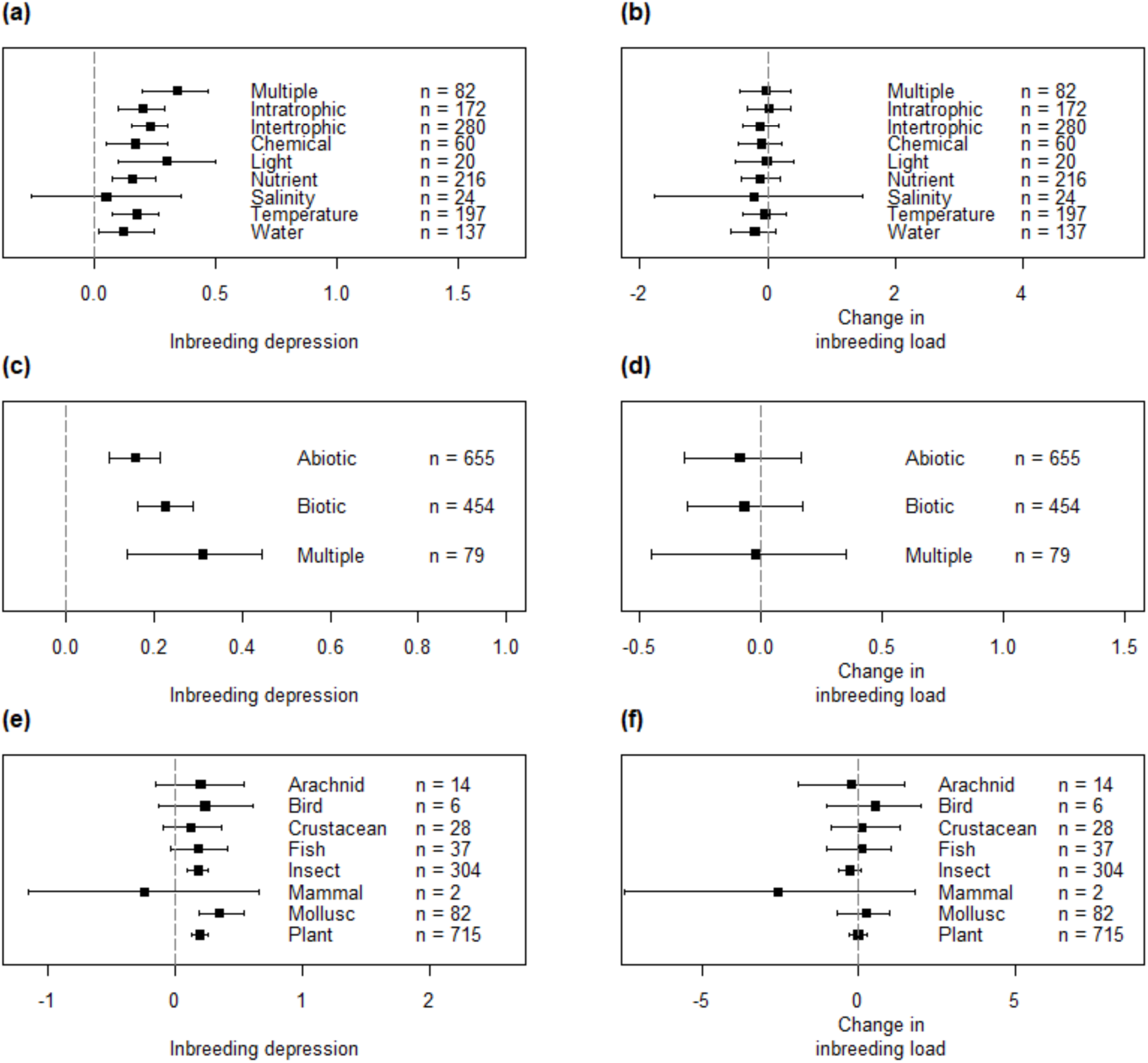
Variation in inbreeding depression or stress-induced change in inbreeding load due to inbreeding among environmental factors and taxonomic groups (using data for all traits). **(a)** inbreeding depression in different stressful environments. **(b)** stress-induced changes in inbreeding load in different environments. **(c)** magnitude of inbreeding depression under different high-level environmental stress categories. **(d)** stress-induced change in inbreeding load in different high-level environmental categories. **(e)** inbreeding depression in different taxonomic groups in stressful environments. **(f)** stress-induced change in inbreeding load in different taxonomic groups. Positive values indicate the expression of inbreeding depression ((a), (c), (e)), or greater inbreeding depression in stressful relative to benign environments ((b), (d), (f)). Effects shown in (a), (c), (e) were estimated at an inbreeding coefficient of 0.5, whereas estimates of changes in genetic load shown in (b), (d), (f) are standardised by the inbreeding coefficient (see Methods). Error bars show 95% credible intervals.

### 2.3 Other sources of heterogeneity

Taxonomic group was a weak and non-significant predictor of both inbreeding depression (Fig. 3e) and changes in inbreeding load between benign and stressful environments (Fig. 3f). However, between- and within-article variation in inbreeding depression in fitness component traits was substantial (47.62% and 46.26% of total variance, respectively; *SI Appendix*; Fig. S7), with a similar pattern for stress-induced shifts in inbreeding load (74.12% and 21.65% of total variation). In contrast, for non-fitness component traits, heterogeneity was dominated by within-article variation, which explained 89.65% of the variation in inbreeding depression. Within-article variation in these non-fitness traits accounted for 32.37% of total variance for stress-induced changes in load.

## 3 DISCUSSION

Our aim was to understand how inbreeding depression is modified by environmental stress. Our meta-analysis revealed widespread and significant inbreeding depression across taxa and environments, and exposed environment-dependent inbreeding depression. Overall—and in contrast to previous evidence syntheses—inbreeding depression was significantly greater in benign than in stressful environments. For fitness components (survival, viability, fecundity, fitness), we detected a unimodal (humped) relationship between the intensity of stress and both the magnitude of inbreeding depression and the shift in inbreeding load between benign and stressful environments, implying that inbreeding depression in stressful environments peaks at intermediate stress levels and declines under extreme stress. For non-fitness component traits, there was a weakly supported U-shaped relationship between shifts in inbreeding load and stress intensity, but no relationship between stress intensity and inbreeding depression.

### 3.1 Environment-dependent inbreeding depression in fitness components

Our findings—that both inbreeding depression and stress-induced changes in inbreeding load have a unimodal relationship with environmental stress—overturns our previous understanding that changes in inbreeding depression should scale positively and linearly with stress intensity (14, 27). Indeed, within populations, we found that in highly stressful environments, inbreeding depression was consistently lower than in benign environments, whilst for lower levels of environmental stress there was no consistent change in inbreeding load between benign and stressful environments. The differences between earlier evidence syntheses and our own could stem from large differences in statistical power and study scope. The Fox & Reed meta-analysis (14) synthesised the impacts of environmental stress on inbreeding depression by comparing the difference in the number of lethal equivalents expressed in stressful versus benign environments within a population (equivalent to the effect size measure we used). However, their meta-analysis only comprised 33 articles with 60 effect sizes, compared with our meta-analysis, which had 116 articles and 2127 effect sizes. Furthermore, in our meta-analysis we used systematic methods for assessing study relevance ((14); Methods section 2.4). Additionally, Fox & Reed’s (14) review was restricted to a single fitness trait (survival). Other fitness components, which are important determinants of population dynamics, were not considered. Using only data for survival in our model, there was still no evidence for a positive linear relationship between changes in inbreeding load and environmental stress. Instead, we detected a unimodal (humped) relationship (*P* = 0.054, *SI Appendix*; Fig. S8), a result consistent with our combined analysis of fitness component traits. Consequently, our study better captures the pattern of co-variation between environmental stress and inbreeding depression, and provides a more general view.

Since individuals cannot have fitness values of less than zero, inbreeding depression should decline under increasingly intense environmental stress towards zero, implying a non-linear relationship between environment-dependent inbreeding depression and environmental stress (28–30). Springer and colleagues (29) found that although environmental stress could increase inbreeding depression in cowpea seed beetles (*Callosobruchus maculatus*), this interaction was weak when environmental stress was very strong. Earlier studies on the same species revealed (i) limited evidence for interactions between inbreeding depression and environmental stress when stress was weak (31) and (ii) clear evidence for increases in inbreeding depression under intermediate environmental stress (14). Hence, Springer et al. (29) concluded that the rate of increase of inbreeding depression with environmental stress decreases when stress itself becomes extreme. Similarly, studying nutritional stress in *Drosophila*, Schou and co-workers found a smaller increase in inbreeding depression when stress intensity was intermediate (28), with inbreeding depression reaching an asymptote. However, neither Schou et al. (28) nor Springer et al. (29) presented evidence for a descending slope in the inbreeding depression-environmental stress relationship. In our meta-analysis, we found that in stressful environments (i) the strongest inbreeding depression occurred when stress levels were intermediate and (ii) inbreeding depression diminished when stress levels increased from intermediate to high values (Fig. 2a). Under severe stress, inbreeding load in benign environments was also consistently greater than that in stressful environments (Fig. 2c). However, excluding cases with a stress intensity exceeding 0.75 eliminated the second-order polynomial pattern, implying that this relationship was driven by extreme stress-linked fitness reductions that constrained space for inbreeding depression. (*SI Appendix*; Fig. S9). By synthesising findings from 116 papers, our meta-analysis provides strong evidence for a non-linear relationship between inbreeding depression and environmental stress levels. Our findings imply that when stress is intense, and fitness decreases towards zero, environmental stress is the main concern for fitness and therefore population viability (29), and the relative importance of inbreeding depression declines.

### 3.2 Environment-dependent inbreeding depression in non-fitness traits

For non-fitness traits, our meta-analysis detected only non-significant, or weakly supported relationships between stress levels and the expected (average) level of inbreeding depression (Fig. 2b) or stress-induced changes in inbreeding load (relative to benign environments; Fig. 2d). These observations might reflect dissimilarities in genetic architecture between fitness components and other traits which modify both the expected costs of inbreeding and the magnitude of shifts in inbreeding depression induced by stress. Components of fitness are thought to be characterised by directional dominance, underpinned by deleterious recessive variants (other dominant mutations that affect fitness are either rapidly fixed or lost). Quantitative (morphological) traits—which were key non-fitness components in our study—often undergo stabilising selection for an intermediate optimum, resulting in weakly directional dominance (32–34), with deleterious mutations that could alter the mean phenotype in either direction. Therefore, inbreeding depression is likely to be greater in fitness components than in other traits (35), as we found in our analysis. The lower levels of inbreeding depression in non-fitness component traits could, in turn, lower the power to detect a consistent shift in inbreeding depression with environmental stress. Similarly, if the phenotype is decanalised by stress (36), exposing novel genetic variation, the exposed variants are more likely to have bidirectional phenotypic effects in quantitative traits, muting phenotypic shifts under inbreeding.

A second possibility is that stress can modify the direction of the relationship between phenotypes and fitness in non-fitness traits (28, 37). For example, plant biomass production might scale positively with fitness in a benign environment, but negatively under drought stress. This would complicate the interpretation of inbreeding depression in non-fitness traits during a shift between benign and stressful environments. In our study, it was necessary to assume a direction for the phenotype-fitness relationship for non-fitness traits, based on arguments and evidence presented in research articles. Stress-induced changes in the phenotype-fitness mapping would undermine this assumption.

### 3.3 Heterogeneity in inbreeding depression

We detected significant heterogeneity in inbreeding depression and stress-induced changes in in genetic load both between and within articles, reflecting species-, or population-specific effects, trait choices and study design. Earlier reviews have shown that species characteristics (e.g. mating system and lifespan) can shape the magnitude of inbreeding depression, generating heterogeneity across taxa (30, 38–40). However, we did not find significant variation in stress-induced changes in inbreeding depression among taxonomic groups, suggesting broadly similar patterns of environment-dependent inbreeding depression across taxa. Neither did we find evidence of consistent differences in inbreeding depression between different types of environmental stress. Many stressors are known to induce highly specific physiological responses (41, 42), which drive stress-specific changes in the fitness effects of recessive deleterious mutations, i.e. conditionally deleterious mutations (18, 24). In this scenario, purging may also amplify differences in the expression of inbreeding depression if a population has had prior exposure to some environmental stresses but not others (2, 33). The lack of taxonomic and environmental variation in inbreeding responses we observed likely reflects wide heterogeneity in species- or population-level inbreeding responses.

### 3.4 Conservation implications

The effective integration and management of genetic issues in conservation practice requires consensus over the available evidence (4, 43). While the relationship between environmental stress and inbreeding depression is a critical issue in conservation, wide heterogeneity in the outcome of studies in experiments and in the wild (37, 44, 45) has, until now, precluded consensus and translation into conservation practice. By comprehensively synthesising data across studies and taxa our meta-analysis provides the synthetic understanding necessary to inform conservation.

Our results imply that while costs to fitness from inbreeding are ubiquitous, and significant, a consistent environmental shift in these costs only occurs under highly stressful conditions. Under these circumstances, inbreeding depression is lower than in benign environments, and the direct effects of environmental stress on fitness therefore strongly outweigh those accruing from inbreeding depression. Nonetheless, for a minority of populations, inbreeding-environment interactions that reduce fitness could occur in the wild—especially at intermediate stress levels (Fig. 2a). In these cases, routine conservation practice that alleviates environmental stress may also reduce the extent of inbreeding depression. The heterogeneity in inbreeding depression-environment interactions we observed does not imply that inbreeding depression is not a concern in conservation practice. Indeed, given the ubiquitous and strong nature of inbreeding depression, the genetic rescue of inbred populations through managed migration is, and should remain, an important conservation tool. This will require the careful management of connectivity between populations (especially where populations were recently connected), and the targeted genetic rescue of populations in which the costs of inbreeding are likely to limit demographic sustainability (6, 46).

### 3.5 Conclusion

Our comprehensive meta-analysis resolves uncertainty in the relationship between inbreeding depression and environmental stress. Specifically, our work confirms the existence of environment-dependent inbreeding depression but overturns the longstanding idea that inbreeding depression is more intense in stressful environments. Furthermore, it shows that, as hypothesised, the relationship between inbreeding depression and environmental stress intensity is unimodal for fitness component traits, not linear, revealing the conditions under which inbreeding depression and environmental stress shape fitness interactively. Thus, while conservation practitioners should continue to manage the deleterious impacts of inbreeding in populations, they should not assume that stressful environments necessarily exacerbate inbreeding depression.

## 4 MATERIALS AND METHODS

### 4.1 Literature search

We used a comprehensive systematic literature map of the fitness and phenotypic consequences of inbreeding depression compiled by Neaves et al. (47), comprising a literature database supporting meta-data and a fully documented review protocol compatible with systematic review methodology. We carried out a literature search to update this resource on 4 May 2018 using improved literature search strings designed by Neaves *et al*. (*SI Appendix*; Table S1 31). Literature searches were conducted in Web of sciences (core collection), Scopus, and JSTOR.

The relevance of articles in the combined literature database was evaluated hierarchically, through sequential assessment of article titles, abstracts and full-texts against our inclusion and exclusion criteria (*SI Appendix;* Fig S1). Briefly, we included articles that recorded the phenotypic and fitness difference between inbred and outbred offspring under contrasting environmental conditions, encompassing experimental manipulations (99.75% of relevant articles), reciprocal transplant experiments (0.17%), and field studies (0.08%). Articles were deemed relevant only when study populations were natural or had been derived from natural populations, with no history of artificial selection.

### 4.2 Data extraction

From each article, we extracted (ⅰ) mean phenotypic or fitness, sample sizes and standard deviation sufficient to calculate inbreeding depression effect sizes in different environmental settings, (ⅱ) the type of environment in which inbreeding responses were observed, and (ⅲ) other effect modifiers that may predict variation in inbreeding depression (e.g. inbreeding coefficient, taxonomic and trait-related information). Typically, this involved extraction of phenotypic data for non-inbred (outbred) and inbred groups of individuals in at least two environments. Binomial traits (e.g., hatched offspring, survival) were extracted as a proportion. If not declared, the standard deviation for binomial data was calculated as:

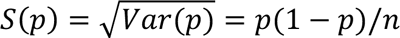

Where p is the probability of occurrence in n observations (48). WebPlotDigitizer version 4.4 (49) was used to extract trait means and standard deviation from figures. Our data followed a nested structure where multiple inbreeding depression effect sizes (“studies’) could be extracted from each relevant publication (“article”). For example, different Study IDs were used for different traits, environmental contrasts, or different levels of inbreeding within an article. We approached the authors to request data if not presented in a usable form.

The environmental manipulations applied to, or experienced by, inbred and outbred groups are listed in Table S2. The benign environment was identified as that environment in which a fitness-component trait (survival; fecundity, fitness), or a trait with known positive relationship to fitness, was greatest in non-inbred individuals. Comparator environments with lower fitness/ phenotypic performance in non-inbred individuals were defined as stressful. Environments declared by the authors of articles as stressful or non-control qualified as stressful in our meta-analysis in 78.1% of cases. For environments that were species interactions, we categorized any interactions over different trophic levels as intertrophic (e.g. predators, disease) and interactions within the same trophic level as intratrophic (e.g. parental care, competition). Environments in which more than one environmental treatment was applied simultaneously or where the environment involved multiple environmental factors varied were categorized as “multiple factors”.

### 4.3 Effect sizes and measurement error variance

Inbreeding effect size, measuring the strength of inbreeding depression, was calculated for each study as a log response ratio:

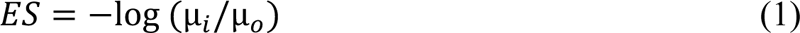

where µ_*i*_ and µ_*o*_ are mean phenotypes for the inbred and non-inbred groups respectively (50). The study measurement error variance (*mev*) was calculated as

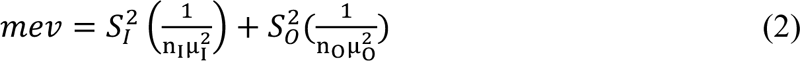

where n_I_ and n_O_ are the sample sizes of inbred offspring and non-inbred offspring and *S*_1_ and *S*_0_ are standard deviations of the inbred and non-inbred groups, respectively (50). Positive values of *ES* indicate the presence of inbreeding depression.

A second effect size was used to investigate population-level changes in inbreeding load between benign and stressful environments, to control for between-study (trait-, population-or species-specific) variation in inbreeding depression in benign environments (environmental response effect size*, ΔES_ENV_*):

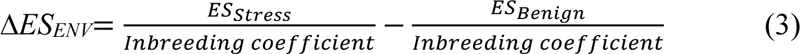

where *Es_stress_* and *Es_benign_* are the inbreeding effect sizes (equation (1)) in the stressful and benign environments. Each of these component effect sizes was standardised by the inbreeding coefficient, rendering the terms 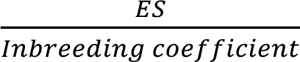 measures of genetic load (51). The associated *mev* was calculated as:

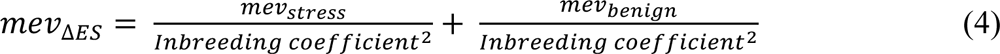

where *mev_stress_* and *mev_benign_* were the original *mev* in their corresponding environments, correcting for the rescaling of the effect size in equation (3).

Stress intensity level was estimated from the mean phenotypes of non-inbred groups and was calculated as:

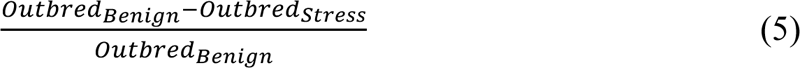

where *Outbred_Benign_* and *Outbred_stress_* are the mean phenotypes of the non-inbred groups in the benign and stressful environments, respectively (14). Measured in this way, stress levels are bound in the interval (0,1) and represent the relative change in phenotype (or fitness) due to environmental change.

### 4.4 Meta-Analysis

Effect sizes were analysed using Bayesian mixed-effects meta-analyses in MCMC_GLMM_ (version 2.29) in R (52, 53). We fitted explanatory variables of interest as fixed effects, including inbreeding coefficient, environmental factors, and stress level. Article ID and *mev* were fitted as random effects in all models to capture and account for variation among and within articles and effect size precision (54). Additional analyses included taxonomic group, trait type, and mating system as fixed effects to understand contextual variation in *ES* and *ΔES_Env_* (*SI Appendix*; Table S4, S5). All analyses of *ES* included the inbreeding coefficient as a fixed effect to control for the variation in the degree of inbreeding used in different studies (centred to allow prediction at an inbreeding coefficient of 0.5). Models assessing the effects of stress intensity on *ES* utilised only effect sizes from stressful environments, since those in benign environments all have a stress value of zero. Best fitting models were identified as those with the lowest DIC.

In our models, there is the potential for an “intrinsic” correlation between the response (effect size) and the stress intensity predictor, with a positive sign (14). This arises because both effect sizes and stress levels share terms in common (i.e. Outbred_stress_). We conducted a numerical simulation to evaluate the direction and strength of this intrinsic relationship given the properties and structure of our data (*SI Appendix*). In our simulation, and following Fox and Reed (14), values for the inbreeding load were drawn independently across environments within studies, allowing us to isolate the effects of the intrinsic correlation (*SI Appendix*; Fig. S2). Analysis of simulated datasets showed that the intrinsic correlation is close to zero for fitness and non-fitness component traits (for *ES*: -0.00317, -0.00324; for Δ*ES_ENV_*: -0.00079, 0.00869), and was not statistically different from zero (*SI Appendix*; Fig. S3).

### 4.5 Publication bias

We tested for publication bias using two regression-based methods that tested the association between effect sizes and their standard errors or effective sample sizes, and by visual inspection of funnel plots (*SI Appendix*; Fig. S4; 55, 56). Since both tests for bias were statistically supported, we provide alternative meta-effect estimates for Figure 1 that are adjusted for publication bias (*SI Appendix*; Fig. S5). None of our statistical results were altered by accounting for publication bias.

## ACKNOWLEDGEMENTS

YFC was supported by an International Scholarship grant from the University of Liverpool and a Government Scholarship to Study Abroad 2019, Ministry of Education, Taiwan. We thank Jenny Hodgson, David Atkinson, Andrea Betancourt, and Ilik Saccheri for constructive feedback on the draft manuscript and J. Hodgson, D. Atkinson and Stephen J. Cornell for advice on statistical analysis.

## AUTHOR CONTRIBUTIONS

YFC: data curation (lead), formal analysis (lead), investigation (lead), visualisation (lead), and writing – original draft preparation (lead), writing – review and editing (supporting). RW: conceptualisation (lead), methodology (lead), supervision (lead), and project administration (lead), writing – review and editing (lead).

## DATA AVAILABILITY STATEMENT

The data and code supporting this study (57) will be available at the University of Liverpool Research Data Repository (Liverpool Elements), on publication: https://doi.org/10.17638/datacat.liverpool.ac.uk/3095.

