## Supporting Information for "Inbreeding depression is greater in benign than in stressful environments"

Raj Whitlock

**This file includes:**

Supporting text

Figures S1 to S11

Tables S1 to S6

SI References

Supporting Information Text

Inclusion and exclusion criteria

Types of study

Our inclusion criteria targeted empirical studies documenting changes in the costs or phenotypic effects of inbreeding across environmental treatments or gradients. Studies were considered relevant when a “control” group of non-inbred individuals and a “treatment” group of inbred individuals could be identified based on either manipulated or observed matings with known familial relationship, or with molecular pedigree data from which the degree of inbreeding could be inferred. We also required that a relevant phenotypic outcome was recorded for both control and treatment groups in more than one environment, and that these environments were either naturally occurring or experimentally manipulated. Thus, relevant articles reported the phenotypic consequences of inbreeding under different biotic or abiotic conditions (environments). Book sections and conference papers were excluded. Review and meta-analysis articles were not included as relevant, but any potentially relevant articles from their bibliographies were assessed against our inclusion criteria.

Types of subjects

Relevant subjects were defined as populations of plants and animals at any location worldwide that were, or were derived from, natural populations without a history of artificial selection. Natural populations were defined to include native populations or introduced, naturalised populations occupying habitats in the absence of human intervention. We excluded studies documenting inbreeding effects resulting from mating among agricultural cultivars or zoo populations since these populations have often been subjected to human-mediated artificial selection (either knowingly or unknowingly). Experimental populations derived from a natural population and maintained at a large population size with minimal artificial selection were also included, and represented the majority of studies (99.14%). If these populations were derived from natural populations within five generations, we labelled them as short-term *ex situ* (76.86%). Populations maintained *ex situ* in a lab or common garden environment for more than five generations were labelled as long-term *ex-situ* populations (22.28%). The remaining studies were observations within natural populations (i.e. in the wild; *in-situ* populations), which made up the minority of cases (0.86 %). This distinction is important, because these non-wild populations might experience different selection regimes relative to natural populations in the wild, and the duration of this exposure may influence the response to inbreeding. Microorganisms were also excluded from our study since (i) the meaning of inbreeding for microorganisms is different from multicellular eukaryotic organisms due to their mechanisms of reproduction and inheritance and (ii) microorganisms are not usually the subject of conservation management interventions, such as genetic rescue.

Types of phenotypic outcome

Relevant outcomes were phenotypic data from the offspring of inbred and outbred matings or crosses under different environmental conditions or treatments. These included fitness-component traits such as fecundity, viability, survival, and compound fitness measures and traits that are less closely related to fitness, such as body size, growth rate and behaviour. Studies were eliminated if the direction of natural selection acting on a trait could not be established or reasonably inferred, or if the authors of articles did not state the direction of this relationship. For example, the direction of the relationship between fitness and a metabolite’s concentration is impossible to infer in the absence of additional data.

Types of comparators

Effect sizes describing the magnitude and direction of phenotypic inbreeding responses were based on a comparison of mean phenotypic comparisons between inbred and non-inbred groups of individuals, with the inbred group acting as comparator. Inbred and outbred groups were defined on the basis of pedigree information (manipulated matings, observed matings or matings inferred from molecular pedigree data). We did not include studies in which the degree of inbreeding was estimated only via molecular data (i.e. FIS; the deviation between observed and expected heterozygosity). Inbred groups of individuals therefore refer to the progenies stemming from inbreeding between related parents, and non-inbred groups arise from either random mating within populations or outcrossing between them. The outbred groups acted as a reference allowing us to quantify the magnitude of inbreeding depression via changes in mean phenotype. In total, 116 articles were included in this meta-analysis (1-116).

Non-environmental effect modifiers

Taxonomic group

Understanding the association between taxonomy and inbreeding depression is essential for conservation practice since it allows conservation biologists to assess the risks of inbreeding in threatened populations. Variation in inbreeding depression may occur among taxa due to taxonomic variation in the mating system, life history and behaviour via breeding patterns and the maintenance or removal of deleterious alleles by natural selection. Domain-level taxonomic classification was recorded for the studied populations as either animal or plant. Animals were further classified as arachnid, bird, crustacean, fish, insect, mammal, and mollusc.

Trait type

Assessing the relationship between different types of phenotypes and inbreeding depression is essential since the effects of inbreeding depression in different traits are a key determinant of population demography. We classified phenotypic traits in which inbreeding depression had been measured as either behavioural, defence, development, fecundity, growth rate, physiology, size, survival, or viability. Most behavioural traits were related to mating behaviour such as mating effort or copulation duration. Defence traits were measured as an organism’s ability to defend against pathogens, predators, herbivores or extreme manipulated environmental conditions, such as heat shock. Developmental traits included the time at reaching developmental milestones, such as age at flowering in plants or developmental disorders, such as fluctuating asymmetry. Reproductive traits included measures of reproductive effort, success or output; measures of the number or probability of reproductive organisms produced. Fitness metrics were compound measures that integrated multiple fitness component traits into one value. These fitness metrics incorporated more than one fitness component trait. Growth rate measured increase in body size or mass per unit of time. Physiological traits mainly had two categories. One was a measure of body composition, such as protein, fat, nitrogen, carotenoid, and chlorophyll content. The other was a measure of physiological function, such as photosynthetic rate or water use efficiency. Size traits were measures of an organism’s total body size or mass or parts of the body’s mass, volume or length. Survival traits were measures of survival proportions at maturity or after establishment. Viability traits represented survival before morphological maturity or establishment or measures of germination or egg hatching rates. “Other” traits were other non-fitness component traits that did not belong to the categories described above.

Fitness component and non-fitness component traits

The genetic architecture of fitness component traits could differ from that for non-fitness component traits, leading to a difference in the expected phenotypic costs of inbreeding (117, 118). Therefore, we characterised phenotypic responses to inbreeding as either fitness component or non-fitness component traits. Fitness component traits included reproductive traits, fitness, survival and viability. The remaining trait types were indirectly linked to fitness and therefore categorised as non-fitness component traits.

Mating system

Mating systems influence the quantity of standing genetic variation and the exposure of recessive deleterious mutations to purging, and therefore shapes the expression and severity of inbreeding depression. Two classifications of mating system were included in this review. In the first (*mating system 1*), the mating system of animals was classified as either polygamous or monogamous. Plant mating systems were classified based on outcrossing rates measured from molecular markers at the population-level. Outcrossing rates less than 0.2 defined selfing populations whereas outcrossing rates greater than 0.8 defined outcrossing populations (119). Any population with an outcrossing rate between 0.2 and 0.8 was defined as a mixed-mating population. Under the second classification (*mating system 2*), taxa were classified as mixed-mating if their offspring were produced from any mixture of selfing and outcrossing, as occurs in many plants or gastropods. Taxa with separate sexes or with self-incompatibility mechanisms were classified as outcrossing. When these data were not provided, additional literature searches were conducted to retrieve the missing information.

Life expectancy

Life expectancy was recorded for animals and plants separately. Life expectancy of animals was recorded in units of days. When data were not provided, literature searches were conducted to retrieve the relevant information.

Plant growth form

Plant growth form has been associated with plant mating system and therefore may also be associated with the magnitude of inbreeding depression (120-122). For example, annual plants are more likely to have mixed mating systems while perennial plants are frequently outcrossing (119, 123).Plants were classified as either annual, biennial, herbaceous perennial, shrub (smaller woody perennial) or tree (larger woody perennial). Additional literature searches were conducted as needed to retrieve missing data.

Base environment of study

Laboratories often represent a benign environment for growth in comparison with field environments, which are considered stressful. As a result, if the expression of inbreeding depression is environmentally contingent, its strength might differ depending on the base environment in which it was measured. Base environment was recorded for each trait where it was measured, and was classified as laboratory, greenhouse, common garden or field.

**Additional methods for meta-analysis**

The MCMC chains for all models were run with a total of 1.1 × 10^5^ iterations with a burn-in of 1 × 10^4^ iterations and then a following thin interval of 100 iterations. The total sample size for the posterior distribution resulting from this sampling scheme was 1000 for each fitted model. Priors for random effects were uniform improper distributions on their standard deviation (124). Posterior distributions were summarised to derive point estimates of the model parameters (posterior means, estimating pooled effect sizes) and their 95% credible intervals. These meta-effect sizes were considered statistically significant when their 95% credible interval did not overlap with zero, or the point estimate of another parameter. Bayesian p-values, representing the probability of the parameter’s location relative to zero, were estimated by examining the proportion of the posterior distribution overlapping with zero (or with the point estimate for another parameter). A Bayesian measure of model fit was assessed using the deviance information criteria (DIC) (125). DIC estimates model adequacy by measuring model fit in relation to complexity (the effective number of parameters) (125). The model with the lowest DIC was selected as the best fitting model. Due to the variation in DIC arising from separate model runs, each model was run three times and the average DIC values were taken and compared.

We determined whether a range of sources of heterogeneity explained variation in they influence inbreeding depression effect sizes and effect sizes measuring changes in inbreeding load with environmental stress (Table. S4; S5). These included the intensity of environmental stress, taxonomic group, the type of environment generating stress, , trait type, fitness class (whether a trait was a component of fitness or not, (126), whether experimental individuals were observed in natural populations or were derived from the wild and maintained in a lab environment for up to five generations (short-term *ex situ*) or longer than five generations (long-term *ex situ*), organism lifespan, mating system (119), and base environment (whether phenotypes were expressed in the field, common garden, greenhouse or lab). We included environmental type and taxonomic group individually as fixed effects to determine whether these variables explained variation in inbreeding depression responses and in separate models as random effects to understand the amount of variation that they explained.

Numerical simulation

Our effect size for inbreeding depression and the stress intensity metric that we used share terms in common, via non-inbred phenotypic means in benign and stressful environments (127). This means that an association between effect size and stress intensity could occur in the absence of any biological signal. To quantify this intrinsic relationship between environmental stress intensity levels and inbreeding effect sizes, we conducted a numerical simulation. In this simulation, stress levels were biologically independent of the change in inbreeding depression between benign and stressful environments, but sill allowed a mathematical dependence between stress level and changes in inbreeding depression. Simulated data recapitulated the structure and properties of our empirical data in all other respects.

Our empirical phenotypic data comprised two types of trait distribution: binomial and quantitative (lognormal). Binomially distributed traits were common in fitness-component traits (including survival and viability), and very rare in non-fitness component traits (only 9 instances out of 492). Therefore, we simulated trait data from a lognormal distribution for non-fitness traits and from a mixture of binomial and lognormal distributions for fitness-related traits. Proposal distributions for simulation and their parameters are shown in Table S3. We checked the comparability of these proposal distributions and the empirical distributions they were meant to mimic using two-sample Kolmogorov-Smirnov tests (results shown in Table S3).

First, we simulated non-inbred mean phenotypic values in benign environments using the proposal distributions in Table S3 with back-transformation of lognormal traits to match the scale and distribution of empirical values (Figure S2). Non-inbred mean phenotypes were simulated in proportion to the number of lognormal and binomial traits in our empirical dataset and to reach a data size that also matched our empirical data (1015 studies and effect sizes). One mean phenotype for the non-inbred group in a benign environment was simulated per study. Next, we simulated stress levels (one per study) using a truncated gamma distribution (or uniform distributions; Table S3), which were good approximations to the empirical distribution of stress levels. Since stress levels are determined by non-inbred mean phenotypes in benign and stressful environments (127), we combined simulated mean phenotypes for non-inbred individuals in the benign environment with simulated stress levels to determine mean phenotypic values for the non-inbred group under environmental stress.

$${Outbred}_{Benign}-(Stress \times{Outbred}_{Benign})$$

Subsequently, we simulated inbreeding loads using a t-distribution (Table S3). Two independent simulated values were used for each study—one for the benign environment, and one for the stressful environment in each study—to ensure that shifts in inbreeding depression between environments were independent of the stress level. A single value of the inbreeding coefficient was also simulated for each study from the empirical probability of coefficients of 0.125, 0.25, 0.5 and 0.75, matching the dominant experimental design where crossing leads to a common inbreeding coefficient in the benign and stressful environments. Within each environment (benign or stressful), inbred mean phenotypic values were calculated using the following equation (Keller & Waller 2002):

${Ib}_{stress}=-\left( Inbreeding load \times F \right)\times{NIb}_{stress}+{NIb}_{stress}$

Next, inbreeding effect sizes in both benign and stressful environments were computed using the corresponding inbred and outbred mean phenotypic values. This allowed us to investigate the intrinsic relationship between simulated stress levels and inbreeding effect sizes since the expected change in inbreeding depression between benign and stress ful environments from biological causes (via changes in the genetic load) is zero.

Simulated phenotypic values, inbreeding loads, inbreeding coefficients and inbreeding effect sizes were compiled into a simulated data frame. To mirror the structure of the empirical dataset, we added an article ID to the simulated data. Since the average number of effect sizes per article is 5.18 in the empirical dataset, we assigned the same article ID to every five simulated effect sizes.

Then, we incorporated simulated residuals, among-article effects and measurement error variances (MEV) into the data frame. Residuals were simulated for each effect size using a normal distribution with a mean of zero and a standard deviation estimated from our empirical posterior residual variance. Bayesian mixed-effects meta-analyses. A common among-article effect was assigned to all effect sizes sharing the same article ID. These values were also simulated from a normal distribution with a mean of zero and a standard deviation estimated from the posterior among-article variance in empirical models. To simulate MEVs which were approximately lognormally distributed, we first generated log-transformed MEVs from a normal distribution, using the mean and standard deviation of the log-transformed MEVs from the empirical data. We then exponentiated these values to obtain the final MEVs on the original scale. The simulated residual variance and among-article variance were added to the simulated effect sizes to reflect the observed variance.

This entire simulation procedure was repeated 1,000 times, resulting in 1,000 independent simulated datasets. We then fit separate Bayesian mixed-effects models to each of these datasets, analysing fitness traits and non-fitness traits independently.

Figures

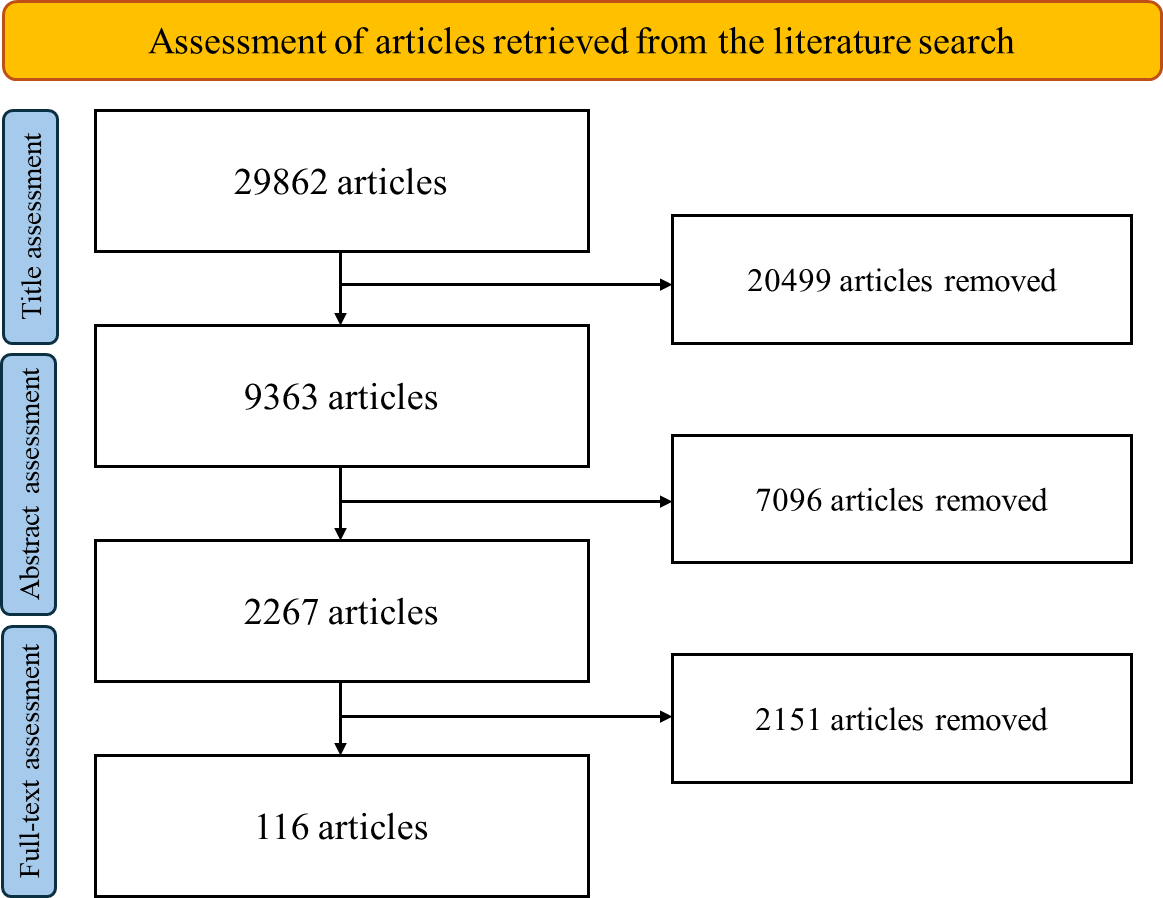

**Fig. S1**. Prisma diagram for the literature relevance assessment. The number in each box indicates the numbers of articles kept or removed after three stages of assessment: title assessment, abstract assessment and full-text assessment.

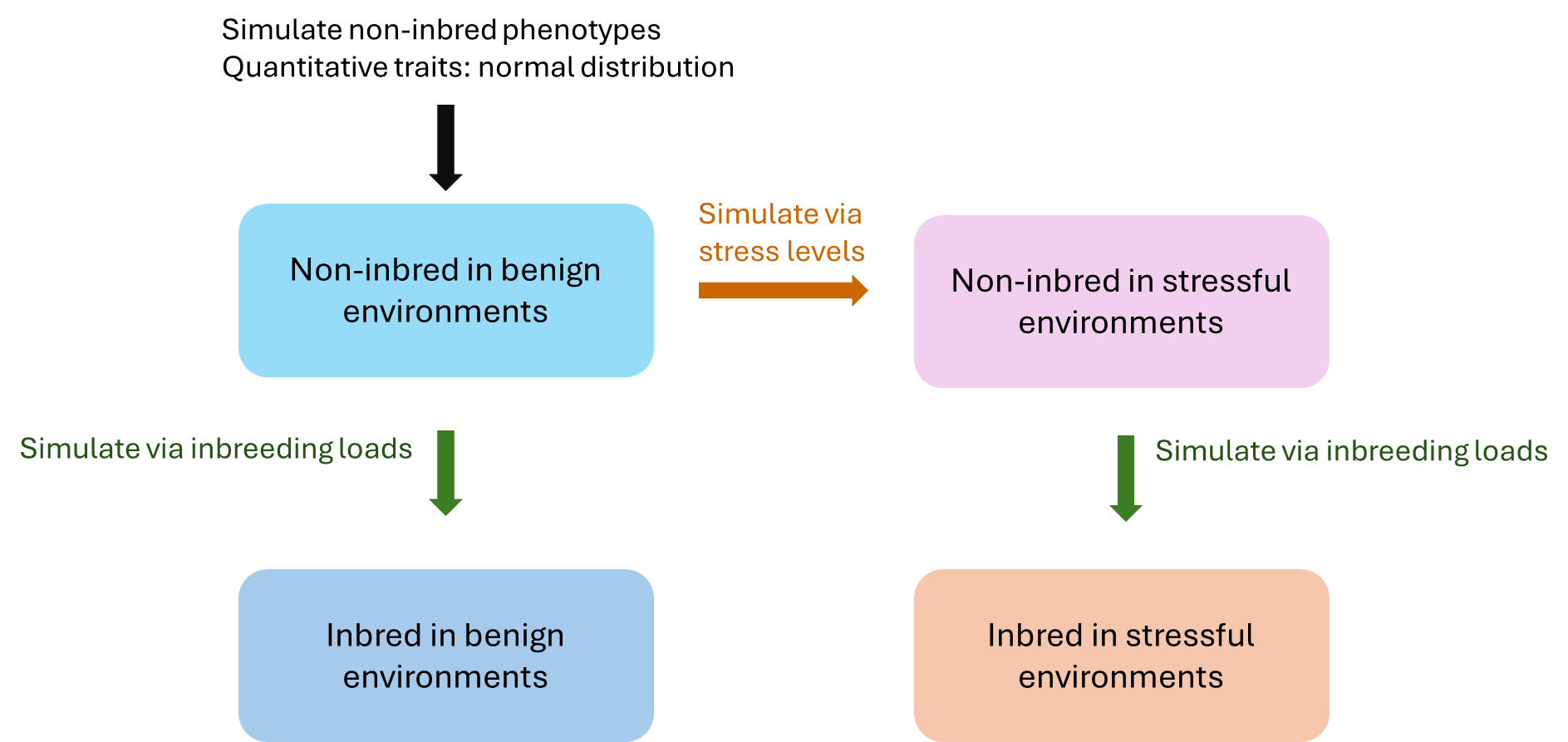

**Fig. S2**. Workflow of numeric simulations. First, non-inbred mean phenotypic values were simulated in benign environments (here, we use quantitative traits as an example). Next, non-inbred mean phenotypic values under environmental stress were simulated using non-inbred mean phenotypes from the benign environment and simulated stress levels. Last, inbred mean phenotypic values under stressful and benign environments were calculated with inbreeding coefficients and inbreeding loads. Two independent draws were made for inbreeding load for each case, rendering the expected change in inbreeding depression to be zero between benign and stressful environments.

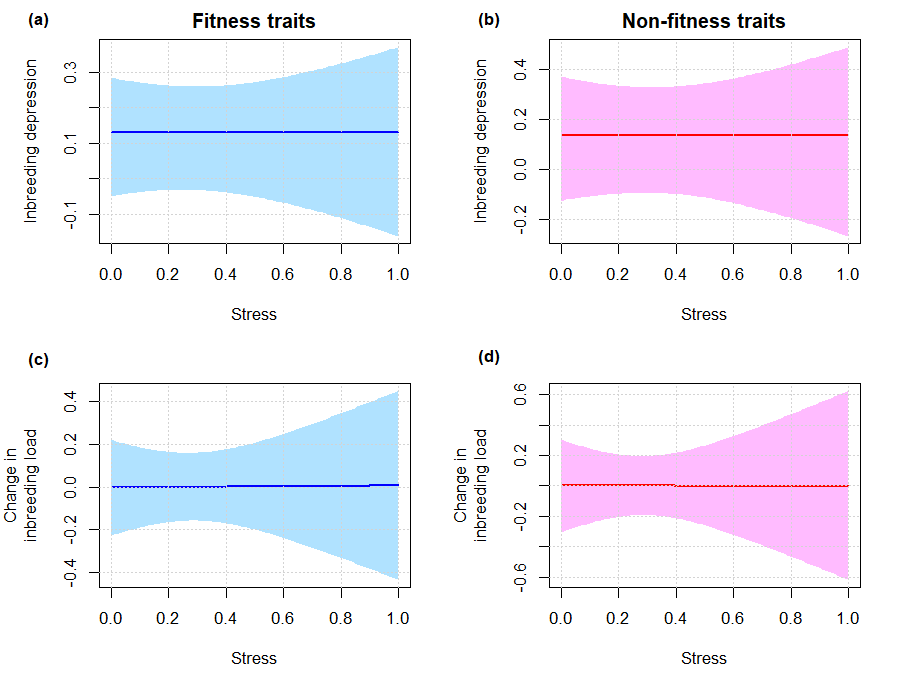

**Fig. S3.** The results of a numeric simulation showing the relationship between environmental stress levels and the expression of inbreeding depression. Each plot shows the best-fitting model prediction for variation of inbreeding depression (a and b) and the change in genetic load due to inbreeding, between benign and stressful environments (c, d) with environmental stress level. The shaded areas represent 95% credible zones for the regression line.

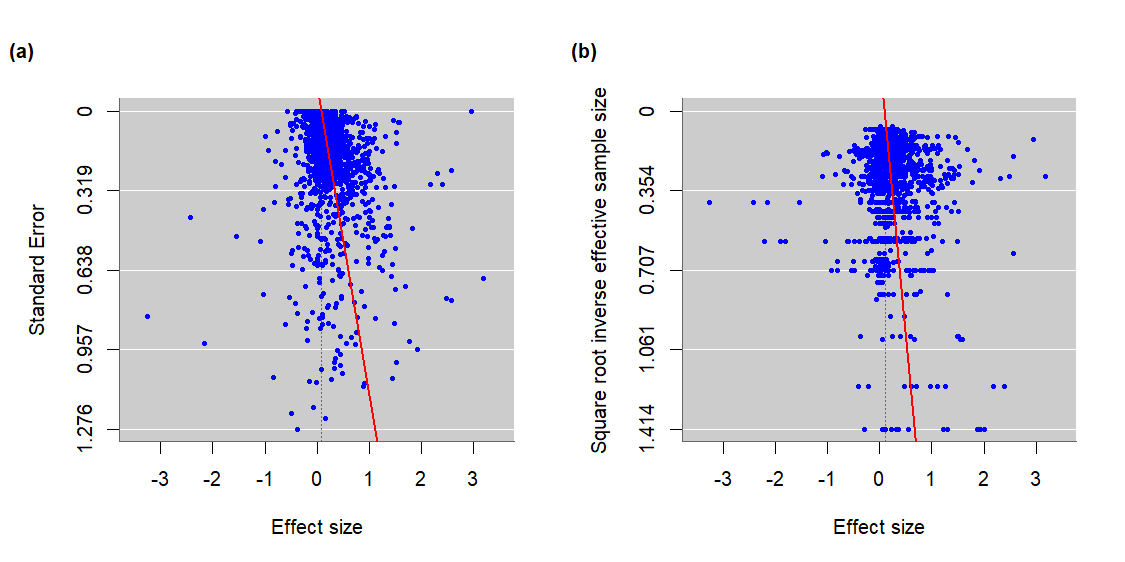

**Fig. S4.** Funnel plots showing the relationship between effect sizes (log response ratio) and (a) their corresponding standard errors; (b) square root inverse effective sample size. Each blue point represents an individual effect size estimate (positive effect sizes indicate inbreeding depression). The red lines indicate the fitted regression used to test for publication bias (variables on the y-axes of these plots were used as the predictors). The slope coefficients are (a) 0.8075, (b) 0.40776 and are both significantly positive (P < 0.0001). This suggests that studies with small sample size or low effect size precision were more likely to publish results when inbreeding depression occurred. For (a), it should be noted that some degree of correlation is expected between log response ratio effect sizes and effect size standard error, due to shared terms in the calculation of these statistics (128). It has been proposed (128) that use of the square-root inverse effective sample size (as in (b)) provides a more robust test for publication bias that avoids this intrinsic relationship between predictor and response.

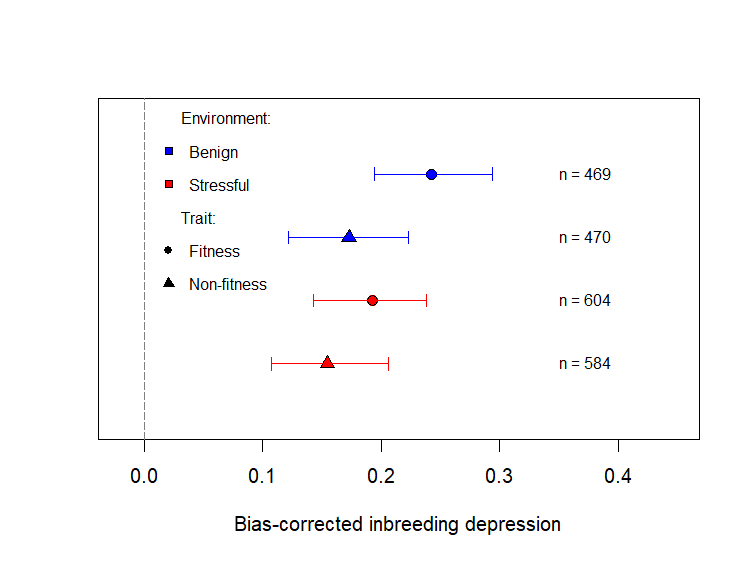

**Fig. S5.** Variation in inbreeding depression between fitness and non-fitness component traits in benign and stressful environments when publication bias is accounted for. Positive values indicate inbreeding depression (a cost to fitness or phenotype). Following (128), the model used here included the inverse of the effective sample size as a predictor, to force parameter estimation at the intercept of this predictor, where effective sample size is infinitely large. Points and error bars (95% credible intervals) show model predictions based on a model that incorporated a trait class by environment interaction (allowing point estimates to vary freely), at an inbreeding coefficient of 0.5. n indicates the number of effect sizes.

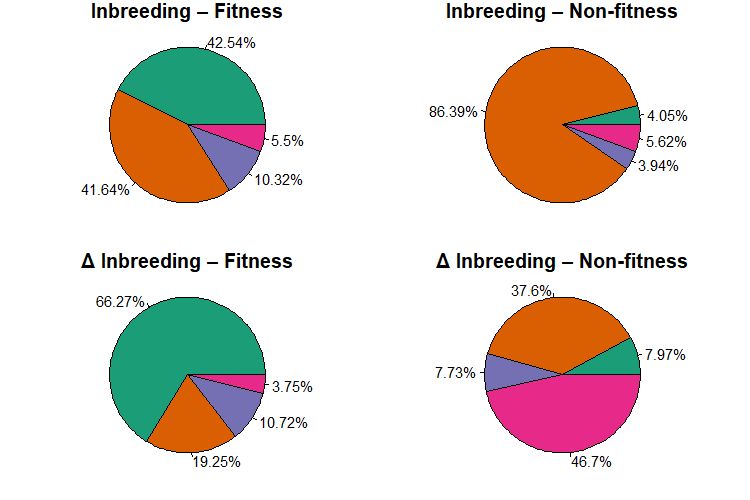

**Fig. S6.** Heterogeneity in inbreeding depression and in stress-induced changes in genetic load, for fitness component and non-fitness traits. Pie charts show the fraction of total variation accounted for by between article variation ■; within article variation ■; taxonomic group ■; and measurement error variance ■. Estimates are derived from models containing random effects for article and taxonomic group, but no other fixed effects predictors.

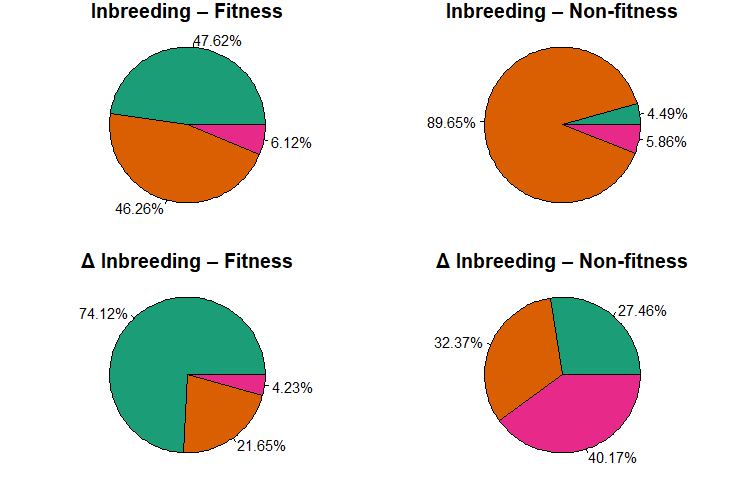

**Fig. S7.** Heterogeneity in inbreeding depression and in stress-induced changes in genetic load, for fitness component and non-fitness traits. Pie charts show the fraction of total variation accounted for by between article variation ■; within article variation ■; and measurement error variance ■. Estimates are derived from models containing random effects for article, but no other fixed effects predictors.

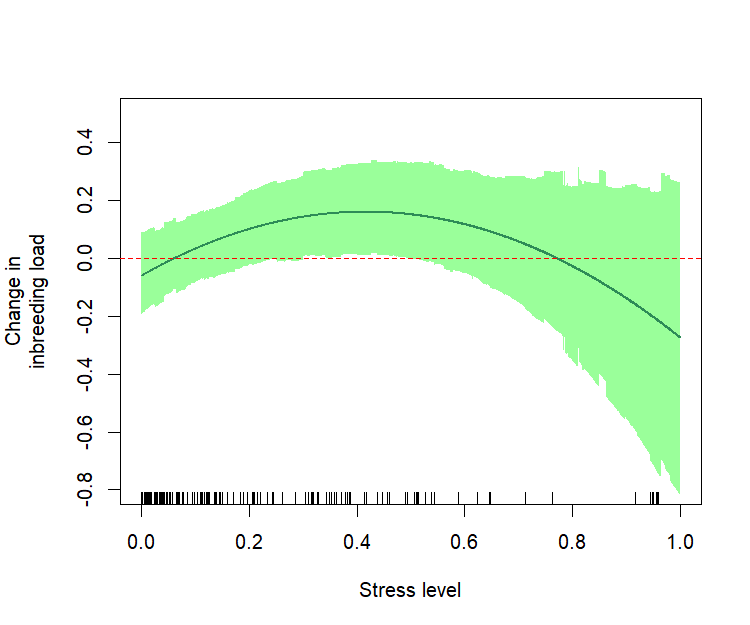

**Fig. S8.** Relationships between environmental stress levels and the change in genetic load between benign and stressful environments for survival (i.e. lethal equivalents; second-order polynomial effect shown; *N* = 181 effect sizes). The plot shows the best-fitting model prediction for variation of change in genetic load with environmental stress level. Stress level data are shown as an x-axis rug. The shaded area represents the 95% credible zone for the regression line based on the posterior distribution.

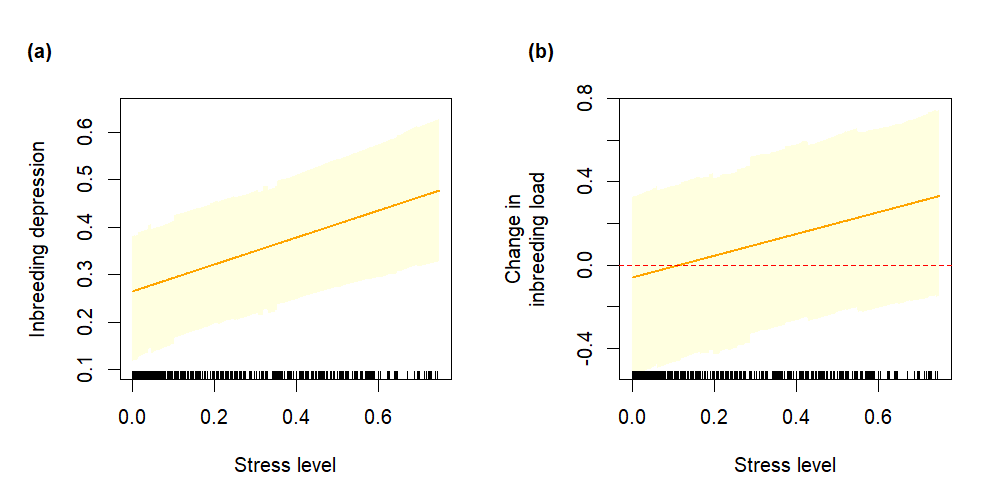

**Fig. S9.** Relationships between environmental stress levels and the expression of inbreeding depression in fitness component traits when stress levels are below 0.75 (i.e. omitting cases with greater stress levels than this). Each plot shows model predictions for variation of inbreeding depression (a) or the change in genetic load between benign and stressful environments (b), with environmental stress level. Stress level data are shown as an x-axis rug (n = 544).

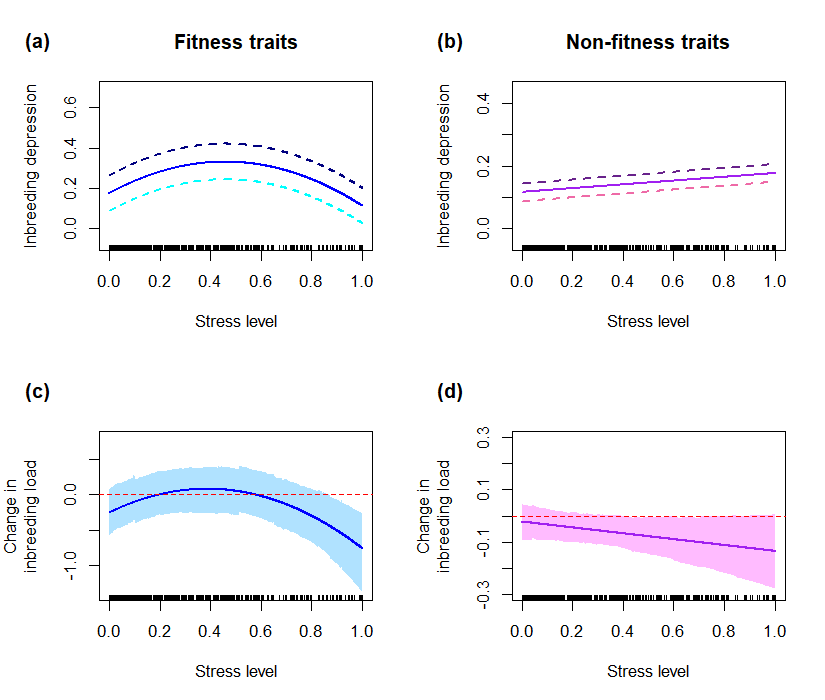

**Fig. S10.** Relationships between environmental stress levels and the expression of inbreeding depression. Each plot shows model predictions for variation of inbreeding depression (a) and (b) or the change in genetic load between benign and stressful environments (c), (d) with environmental stress level. Stress level data are shown as an x-axis rug. (a) the relationship between inbreeding effect sizes and stress level in stressful environments in fitness component traits (second-order polynomial effect shown; N = 602 effect sizes). (b) the relationship be-tween inbreeding effect sizes and stress level in stressful environments in non-fitness component traits (linear effect shown; N = 584). (c) relationship between stress-induced changes in genetic load and stress levels for fitness component traits (second-order polynomial effect shown; N = 602). (d) relationship between stress-induced changes in genetic load and stress levels for non-fitness traits (linear effect shown; N = 584). In (a) and (b), the solid lines indicate inbreeding coefficients at 0.5, whereas the dark/light dash lines indicate inbreeding coefficients at 0.75/0.25. The shaded areas represent 95% credible zones for the regression line in (c) and (d). The red dashed line is y = 0 in (c) and (d).

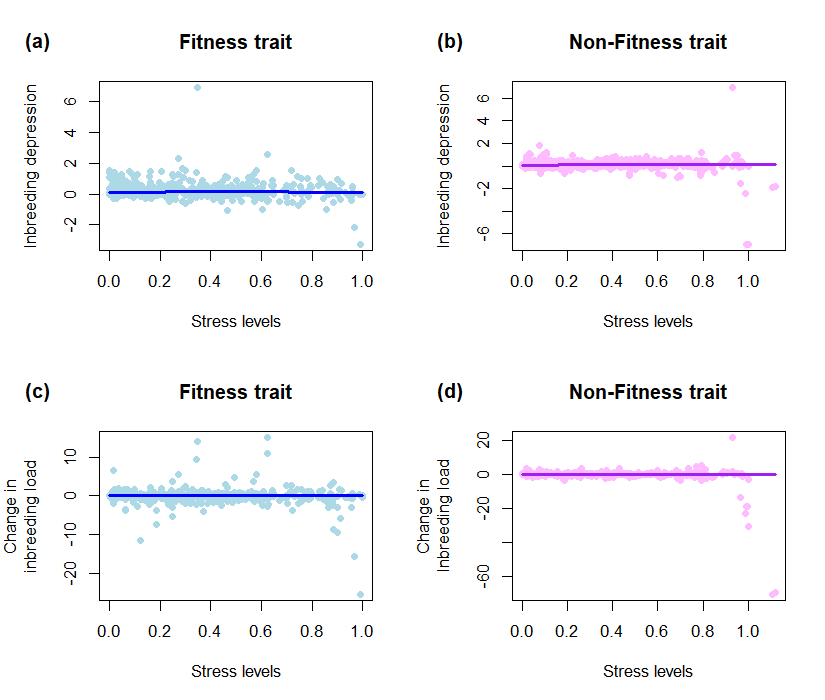

**Fig. S11.** Relationship between stress levels and the expression of inbreeding depression. The empirical data was explored with a generalised additive model (GAM) fit to explore whether linear or polynomial stress predictors could capture the empirical relationship between stress and the expression of inbreeding depression. The points in each plot show the distribution of empirical data for inbreeding depression (a) and (b) or the change in inbreeding load between stressful and benign environments (c) and (d). The lines indicate the model predictions from GAM models.

Tables

**Table S1.** The final search strings used in this meta-analysis to retrieve articles from online database.

| Group | Search string |
| --- | --- |
| Inbreeding related strings | "In*breeding coefficient$" |
|  | "Cost$ NEAR/5 in*breeding" |
|  | (inbred SAME mating*) NOT (("Quantitative trait loc*")OR(QTL*)) |
|  | (inbred SAME (offspring OR progeny)) |
|  | Selfed SAME out* |
|  | "Optimal outcrossing" OR "Outcrossing distance" |
|  | "Benefit* NEAR/5 dispersal" |
|  | "Cost* NEAR/5 dispersal" |
|  | ("Natal dispersal" AND (inbred OR in*breeding OR heterosis OR self* OR fitness)) |
|  | (Philopatr* AND (inbred OR in*breeding OR heterosis OR self* OR fitness)) |
| Fitness related strings | (Depression SAME in*bre*) |
|  | (Depression SAME fitness) |
|  | (Heterosis AND in*breeding) |
|  | "Genetic load" |
| Effects of inbreeding related strings | Effect$ NEAR/5 in*breeding |
|  | consequence$ NEAR/5 Inbreeding |
|  | influence$ NEAR/5 inbreeding |
|  | Outcome$ NEAR/5 inbreeding |

The Boolean syntax above is based on Web of sciences template, and changes were made to adapt the search to the other two literature databases used here.

**Table S2.** Categories of environmental factors used in this meta-analysis.

| Environmental factors that were modified to cause stress | Type of environmental change | Example |
| --- | --- | --- |
| Light | Abiotic | Sunlight availability |
| Nutrient | Abiotic | Food availability |
| Water | Abiotic | Drought |
| Temperature | Abiotic | Increased temperature |
| Chemical | Abiotic | Methanol treatment |
| Salinity | Abiotic | Salt concentration |
| Ventilation | Abiotic | Modified air circulation |
| Intertrophic | Biotic | Pathogen, predators |
| Intratrophic | Biotic | Maternal care, competition |
| Multiple factors | Multiple factors | Food and drought |

**Table S3.** Variables and the proposal distribution for numeric simulations. The R functions used are in parentheses.

| Variables | Distribution for simulation | Kolmogorov-Smirnov p value |
| --- | --- | --- |
| Fitness quantitative trait | Log-normal distribution (rlnorm) | 0.0274 |
| Fitness binomial traits | Gamma distribution (rgamma) | 0.0112 |
| Non-fitness quantitative trait | Normal distribution (rnorm) | 0.4684 |
| Stress levels | Truncated gamma (rtrunc) | 0.0002 |
| Inbreeding loads | t-distribution (rt) | 0.0003 |

The *P* values are the results from the comparisons of empirical and simulated distributions using Kolmogorov-Smirnov test.

**Table S4.** Model fitting summaries for meta-analysis containing fix-effect explanatory variables and inbreeding effect sizes. Bold text indicates best fit models judged by DIC.

| Model | Fixed effects | DIC | Among-article variance | Within-article variance | Within-study variance | Trait types |
| --- | --- | --- | --- | --- | --- | --- |
| A1 | ~Intercept | 265.95 | 0.06665 | 0.06475 | 0.00857 | FC |
| A2 | ~*F* | 251.79 | 0.07138 | 0.06276 | 0.00857 | FC |
| A3 | ~*F* + Stress level | 245.11 | 0.07225 | 0.06191 | 0.00857 | FC |
| **A4** | **~*F* + poly (Stress level, 2)** | **234.15** | **0.07307** | **0.06068** | **0.00857** | **FC** |
| A5 | ~*F* + poly (Stress level, 3) | 231.06 | 0.07127 | 0.0601 | 0.00857 | FC |
| **B1** | **~Intercept** | **660.48** | **0.00769** | **0.15367** | **0.01004** | **NFC** |
| B2 | ~*F* | 662.16 | 0.00809 | 0.154 | 0.01004 | NFC |
| B3 | ~*F* + Stress level | 662.08 | 0.00733 | 0.1542 | 0.01004 | NFC |
| B4 | ~F + poly (Stress level, 2) | 665.07 | 0.00729 | 0.1547 | 0.01004 | NFC |
| E1 | ~Intercept | 1088.44 | 0.03145 | 0.1159 | 0.00954 | All |
| **E2** | **~*F*** | **1082.13** | **0.03495** | **0.1149** | **0.00954** | **All** |
| E3 | ~*F* + Env_factors | 1093.98 | 0.02906 | 0.1163 | 0.00954 | All |
| E4 | ~*F* + Type_Env | 1084.23 | 0.03103 | 0.1155 | 0.00954 | All |
| E5 | ~*F* + Taxonomic group | 1086.19 | 0.03625 | 0.1152 | 0.00954 | All |
| E6 | ~*F* + Short-term | 1083.06 | 0.03556 | 0.1155 | 0.00954 | All |
| E7 | ~*F* + Mating_system1 | 1084.48 | 0.03786 | 0.1148 | 0.00954 | All |
| E8 | ~*F* + Mating_system2 | 1082.91 | 0.03461 | 0.115 | 0.00954 | All |
| E9 | ~*F* + Base_Env | 1083.59 | 0.03626 | 0.115 | 0.00954 | All |

*F*: inbreeding coefficient; FC: fitness component traits; NFC: non-fitness component traits; All: all types of traits. Env_factors: environmental factors that were modified to cause stress listed in Table 1;Type_Env: types of environmental change listed in Table 1; Trait: categorial description of the traits, whether they are fitness component traits or not. The details of base environment (Base_Env), mating_system1 and mating system2 were described in “Description of non-environmental effect modifiers”. Whether a population is a short-term ex situ population or not was defined in “Inclusion and exclusion criteria: type of subjects”.

**Table S5.** Model fitting summarises for meta-analysis containing fix-effect explanatory variables and ∆ ES_ENV_. Bold text indicates best fit models judged by DIC.

| Model | Fixed effects | DIC | Among-article variance | Among-study variance | Within-study variance | Trait types |
| --- | --- | --- | --- | --- | --- | --- |
| A1 | ~Intercept | 1507.89 | 1.637 | 0.47827 | 0.09342 | FC |
| A2 | ~Stress level | 1510.31 | 1.63667 | 0.48123 | 0.09342 | FC |
| **A3** | **~poly (Stress level, 2)** | **1497.09** | **1.71467** | **0.46717** | **0.09342** | **FC** |
| A4 | ~poly (Stress level, 3) | 1495.93 | 1.70333 | 0.4662 | 0.09342 | FC |
| B1 | ~Intercept | 582.01 | 0.07765 | 0.09154 | 0.11361 | NFC |
| B2 | ~Stress level | 583.19 | 0.02063 | 0.09147 | 0.11361 | NFC |
| **B3** | **~poly (Stress level, 2)** | **573.71** | **0.02258** | **0.08995** | **0.11361** | **NFC** |
| E1 | ~Intercept | 2375.68 | 1.184 | 0.282 | 0.10724 | All |
| E2 | ~Env_factors | 2391.98 | 1.8015 | 0.11603 | 0.10724 | All |
| E3 | ~Type_Env | 2379.39 | 1.19133 | 0.28287 | 0.10724 | All |
| **E4** | **~Taxonomic group** | **2370.28** | **1.25667** | **0.28063** | **0.10724** | **All** |
| E5 | ~Short_term | 2376.04 | 1.177 | 0.2829 | 0.10724 | All |
| E6 | ~Mating_system1 | 2371.71 | 1.18 | 0.28 | 0.10724 | All |
| E7 | ~Mating_system2 | 2373.65 | 1.179 | 0.2814 | 0.10724 | All |
| E8 | ~Base_Env | 2381.25 | 1.141 | 0.2831 | 0.10724 | All |

*F*: inbreeding coefficient; FC: fitness component traits; NFC: non-fitness component traits; All: all types of traits. Env_factors: environmental factors that were modified to cause stress listed in Table 1; Type_Env: types of environmental change listed in Table 1. The details of base environment (Base_Env), mating_system1 and mating system2 were described in “Description of non-environmental effect modifiers”. Whether a population is a short-term ex situ population or not was defined in “Inclusion and exclusion criteria: type of subjects”.

**Table S6.** Changes in genetic load between benign and stressful environments as measured by the proportion of significantly positive, negative and non-significant environment-response effect sizes in different ranges of environmental stress. Positive effect sizes reveal that genetic load was greater in a stressful than in a benign environment. The statistical significance and direction of each effect size was determined by comparing its confidence interval (derived from the measurement error variance (129) with zero. Values in the table represent the empirical distribution of effects rather than model predictions from meta-analysis. However, they also do not represent a formal meta-analysis as they do not make an estimate of the mean effect that incorporates a weighting by effect size precision.

| **Stress level** | **Significantly negative** | **Significantly positive** | **Non-significant** | **Fitness class** |
| --- | --- | --- | --- | --- |
| 0–0.35 | 16.45% | 13.26% | 70.29% | Fitness |
| 0.35–0.65 | 20.13% | 20.13% | 59.73% | Fitness |
| 0.65–1 | 23.08% | 8.97% | 67.95% | Fitness |
| 0–0.35 | 7.69% | 6.32% | 85.99% | Non-fitness |
| 0.35–0.65 | 7.46% | 3.73% | 88.81% | Non-fitness |
| 0.65–1 | 13.95% | 9.31% | 76.74% | Non-fitness |
